# Liver Protects Cytoarchitecture, Neuron Viability and Electrocortical Activity in Post-Cardiac Arrest Brain Injury

**DOI:** 10.1101/2023.12.13.571604

**Authors:** Zhiyong Guo, Meixian Yin, Chengjun Sun, Guixing Xu, Tielong Wang, Zehua Jia, Zhiheng Zhang, Caihui Zhu, Donghua Zheng, Linhe Wang, Shanzhou Huang, Di Liu, Yixi Zhang, Rongxing Xie, Ningxin Gao, Ping Yu, Liqiang Zhan, Shujiao He, Yifan Zhu, Yuexin Li, Björn Nashan, Schlegel Andrea, Jin Xu, Qiang Zhao, Xiaoshun He

## Abstract

**BACKGROUND:** Brain injury is the major reason for patient deaths in victims who survive after cardiac arrest. Clinical studies have shown that the presence of hypoxic hepatitis and pre-cardiac arrest liver disease is associated with increased mortality and inferior neurological recovery. However, how the liver might impact the pathogenesis of post-cardiac arrest is still unknown.

**METHODS:** An *in vivo* global cerebral ischemia model was established to assess how simultaneous liver ischemia affected the recovery of brain ischemic injury. In addition, an *ex vivo* brain normothermic machine perfusion (NMP) model was established to evaluate how addition of a functioning liver might impact the circulation, cytoarchitecture, neuron viability and electrocortical activity of the reperfused brain post-cardiac arrest.

**RESULTS:** In the *in vivo* model, we observed a larger infarct area in the frontal lobe, elevated tissue injury scores in the CA1 region, as well as increased intravascular immune cell adhesion in the reperfused brains with hepatic ischemia, compared to those without simultaneous hepatic ischemia. The results of the *ex vivo* model demonstrated that the addition of a functioning liver to the brain NMP circuit significantly reduced post-cardiac arrest brain injury, increased neuronal viability and improved electrocortical activity. Furthermore, we observed significant alterations in gene expressions and metabolites in the presence or absence of hepatic ischemia.

**CONCLUSIONS:** Our research highlights the crucial role of the liver in the pathogenesis of post-cardiac arrest brain injury. These findings shed lights on a cardio-pulmonary-hepatic brain resuscitation strategy for patients with cardiac arrest.

## INTRODUCTION

Sudden cardiac arrest (CA) refers to an unexpected arrest or even death from a cardiovascular cause, which remains a major public health problem estimated to account for 50% of all cardiovascular deaths^1^. Every year estimated 375,000–700,000 citizens are suffering CA in Europe and the USA^2,3^. Although progress has been made in cardiopulmonary resuscitation (CPR) and life support techniques, post-CA survival rates are poor, varying between 8–23%^4–7^.

Mortality after CA is mainly triggered by post-CA shock and brain injury^8^. After return of spontaneous circulation, the whole-body ischemia-reperfusion response results in the post-CA syndrome, comprising post-CA brain injury, post-CA myocardial dysfunction, the systemic ischemia-reperfusion response and persistent precipitating pathology, leading to significant morbidity and mortality^9^. Indeed, multiorgan dysfunction post-CA largely affects the recovery of brain injury^9^. Nevertheless, the roles of individual organs other than the heart (circulation) in post-CA brain injury are almost unknown, limiting the development of novel therapeutic strategies.

Studies have shown that hypoxic hepatitis (HH) occurs in 7-21% post-CA patients and is significantly associated with poor neurological outcome, high mortality rate and long intensive care unit (IUC) stay^10–13^. On the other hand, it has been demonstrated that cardiac arrest survivors with cirrhosis have worse outcome than those without. Particularly, no patients with Child-Turcotte-Pugh C cirrhosis and advanced acute-on-chronic liver failure survived 28 days with good neurological outcome^13^. Similarly, the results from a retrospective observation study, using a nationwide population-based out-of-patient cardiac arrest (OHCA) registry, have also shown that the overall clinical and neurological outcomes are poorer in OHCA patients with liver cirrhosis than those without^14^. Collectively, these clinical data suggest that the liver might play a crucial role in the pathogenesis of post-CA brain injury. However, if the liver is a victim or an offender in post-CA brain injury is unclear.

In the current study, results from the *in vivo* global cerebral ischemia model show that ischemic injuries of the brains with simultaneous liver ischemia are exacerbated compared to those without liver ischemia. And the results of the *ex vivo* brain normothermic machine perfusion (NMP) model show that a well-functioning liver can protect post-CA brain injury. These results of the current study provide direct evidence supporting the crucial role of the liver in the pathogenesis of post-CA brain injury.

## METHODS

Detailed methods are listed in the Supplemental Material. The methods used to implement the research will be made available to any researcher for purposes of reproducing the results. The study protocol was approved by the Animal Ethical Committee of our institute ([2019]193).

### *In vivo* Global Cerebral Ischemia with or without Liver Ischemia

Thirteen Tibet minipigs (25.0 ± 3.1 kg, 6 months, purchased from Guangdong Pearl Biotechnology Co. LTD) were included in this experiment. 5 pigs underwent 30 minutes global cerebral and hepatic ischemia then reperfusion (BLWI-30 group). 5 pigs underwent 30 minutes global cerebral ischemia then reperfusion (BWI-30 group). 3 pigs underwent Sham operation (control group). During operations, the pigs were intubated and anesthetized with inhaled isoflurane (1%-2%) and propofol (100 mg/h). Pancuronium (0.1 mg/kg) and sufentanil (10 μg/kg) were injected before operation for neuromuscular blockage and perioperative analgesia. Pigs were monitored for 24 hours after operation unless they died prematurely. Blood and tissue samples were obtained at scheduled time points. Antibiotics was given per 8 hours perioperatively and body temperature was kept at 37°C using a heating pad.

The pig model of global cerebral ischemia and neurological severity scores has been established^15^. With a 7-8 cm supra-sternal incision, innominate artery, left subclavian artery, both internal mammary arteries, both vertebral arteries, both common carotid arteries were isolated. Right mammary artery and right external jugular vein were catheterized to obtain blood gas samples and administrate drugs. The innominate artery, the left subclavian just distal to the aortic arch, both internal mammary arteries and both distal subclavian arteries were clamped for global brain ischemia. 250 IU/kg heparin was given before ischemia insult. All vascular clamps were removed after 30 minutes brain ischemia. *In Vivo* Optical Spectroscopy (INVOS), applied to monitor cerebral tissue oxygen saturation, ambulatory blood pressure monitoring and neural examination ensured global brain ischemia completely. 30 minutes after reperfusion, protamine was injected and the incision was closed. A 20 Fr drainage tube was placed for mediastinum drainage. Midazolam (0.1mg/kg) was given when postoperative seizures prolonged (>few minutes).

In the BLWI-30 group, the portal vein and hepatic artery were well dissected. The liver underwent simultaneous ischemia and reperfusion with the brain by clamping and unclamping the two vessels.

### Triphenyl-Tetrazolium Chloride (TTC) Staining

After death or after reperfusion for 24 hours, frontal coronal brain slices of the pigs were cut with a thickness of 2 mm. The slices were put into 2% triphenyl-tetrazolium chloride solution (Asegene, G3005) in a dark and were bathed at 37°C for 30 minutes. After washed with PBS for 3 minutes, brain slices were photographed immediately. The white area indicates infarct tissue.

### Tissue Injury Scores

Indicators of brain tissue damage included disappearance of neurites, cell shrinkage, vacuolization or loosening of tissues, and microvascular damage. the injury scores ranged between 0 and 5 points, which represented no, very mild, mild, moderate, severe, and very severe injury, respectively.

### RNA-seq Sample Preparation, Library Construction, and Data Analysis

Total RNA was extracted from the frontal lobes of 10 biological replicates and temporal lobes of 9 biological replicates, followed by the construction of cDNA libraries using a strand-specific RNA-seq protocol. The libraries were subjected to high-throughput sequencing using DNBSeq T7 platforms. RNA-seq reads reads were mapped to the reference genome (susScr11) using alignment algorithms (STAR version 2.7.3a), and gene expression levels were quantified using established bioinformatics pipelines (featureCounts Version 2.0.1). Differential gene expression analysis was performed to identify genes exhibiting significant changes in expression between the experimental conditions using threshold (log2(fold change) >1 and *P* value<0.05). GO Processes analysis was analyzed using the MetaCore.

### *Ex vivo* Brain NMP Technology

Twenty *Tibet minipigs* (30–50 kg, purchased from Southern Medical University, Guangzhou, China) were used during the development of the brain NMP technology. In these preliminary experiments, the technology, including the NMP conditions, perfusate components, methods for brain function assessment, and the appropriate *ex vivo* NMP duration, was optimized. After that, brains from 27 additional pigs were perfused in the formal experiments, and five more brains were harvested immediately after cardiac arrest and used as controls (Sham group) in the pathological assessments. In our *ex vivo* brain NMP system (**Figure 3A**), a whole-blood-based perfusate (**Table S1**) was circulated in a pulsatile arterial line and in a nonpulsatile portal vein line through the oxygenation unit, heater, and filtration unit, after which it perfused the brain (and liver). The filtration unit filtered out small blood clots and impurities, and the heater maintained the temperature of the perfusate at 37°C. Over 10 minutes, the arterial perfusion pressure was gradually increased from 60 mmHg to 90–95 mmHg to achieve a middle cerebral artery (MCA) flow close to the preoperative value. The portal vein perfusion pressure was set at 5–12 mmHg to maintain a flow higher than 500 mL min^-1^.

The brains were reperfused *ex vivo* rapidly or after various intervals. Warm ischemia time (WIT) was defined as the interval between cardiac arrest and brain reperfusion. When a liver was perfused simultaneously, it was reperfused *ex vivo* as soon as harvested, to ensure its viability. There were six experimental groups in the current study: a brain-only control with rapid NMP (BOR group), a liver-assisted brain with rapid NMP (LABR group), and four liver-assisted brain groups in which brain NMP was preceded by 30-240 minutes WIT (LABWI groups) (**Figure S4A**).

### Global Electroencephalograph (EEG) Monitoring of the Brains

EEG was monitored during *ex vivo* brain NMP. Based on the presence of δ (1 point), θ (2 points), α (3 points) and β waves (4 points), the EEG score was calculated hourly by summing the top two points.

### Detection of Brain Tissue and Perfusate

The brain tissues after death or after reperfusion 24 hours *in vivo*, were performed for hematoxylin-eosin (HE) staining, TUNEL staining, Nissl staining, immunohistochemistry, quantitative reverse transcription-polymerase chain reaction (RT-qPCR), RNA sequencing analysis and metabolomics analysis. The brain tissues after 6 hours of *ex vivo* NMP were prepared for HE staining, the transmission electron microscopy (TEM), Nissl staining, immunohistochemistry, and immunofluorescence. The perfusate were prepared for S100-β measurement and metabolomics analysis.

### LC-MS/MS Analysis for Metabolites of *in vivo* Brain Tissues and *ex vivo* Perfusate

The data were analyzed using SAS 9.0 (SAS Institute Inc.). Statistical significance was declared at *P*<0.05. The resultant three-dimensional data involving the peak number, sample name, and normalized peak area were fed into SIMCA 14.1 (Sartorius Stedim Data Analytics AB; Umea, Sweden) for principal component analysis (PCA) and orthogonal projections to latent structures–discriminant analysis (OPLS–DA). PCA showed the distribution of the original data. To obtain an enhanced level of group separation and to better understand the variables responsible for classification, supervised OPLS–DA was applied. This permutation test was conducted to validate the model. On the basis of OPLS–DA, a loading plot was constructed to show the contribution of variables to the differences between the two groups. To refine this analysis, the first principal component of VIP was obtained. If *P*<0.05 and VIP>1, then the variable was defined as a significantly differential metabolite between the groups. Using the Kyoto Encyclopedia of Genes and Genomes (KEGG, http://www.genome.jp/kegg), the metabolic pathways associated with each differential metabolite were acquired.

### Statistical Analysis

All data are presented as a mean ± SEM or median with interquartile range. *T* test and Mann-Whitney test were used for statistical analyses using GraphPad Prism v9.4.0. For comparisons between multiple experimental groups, ordinary 2-way ANOVA followed by the Sidak’s multiple comparison test was used. *P*<0.05 was considered statistically significant.

## RESULTS

### Simultaneous Hepatic Ischemia Aggravates Brain Injuries in Global Cerebral Ischemia Model

To investigate how the liver might affect post-cardiac arrest brain injury, we established an *in vivo* 30 minutes global cerebral ischemia model with (BLWI-30) or without (BWI-30) simultaneous hepatic ischemia (Figure 1A and 1B). The cerebral tissue oxygen saturation (SctO_2_) was steady in the Sham group, but significantly declined after the blood supply to the brain was blocked in the BLWI-30 and BWI-30 groups (**Figure S1A**). The pigs had higher neurological severity scores in the BLWI-30 and BWI-30 groups than in the Sham group, at 6 hours after reperfusion (**Figure S1B**). In addition, the serum aspartate transaminase (AST) and lactate dehydrogenase (LDH) levels at 4 hours after reperfusion were higher (**Figure S1C and S1D**), and liver tissue damage was more severe at 24 hours after reperfusion (**Figure S1E and S1F**) in the BLWI-30 versus BWI-30 group. These data indicates that the global cerebral ischemia model with or without simultaneous hepatic ischemia has been established successfully.

**Figure 1.**
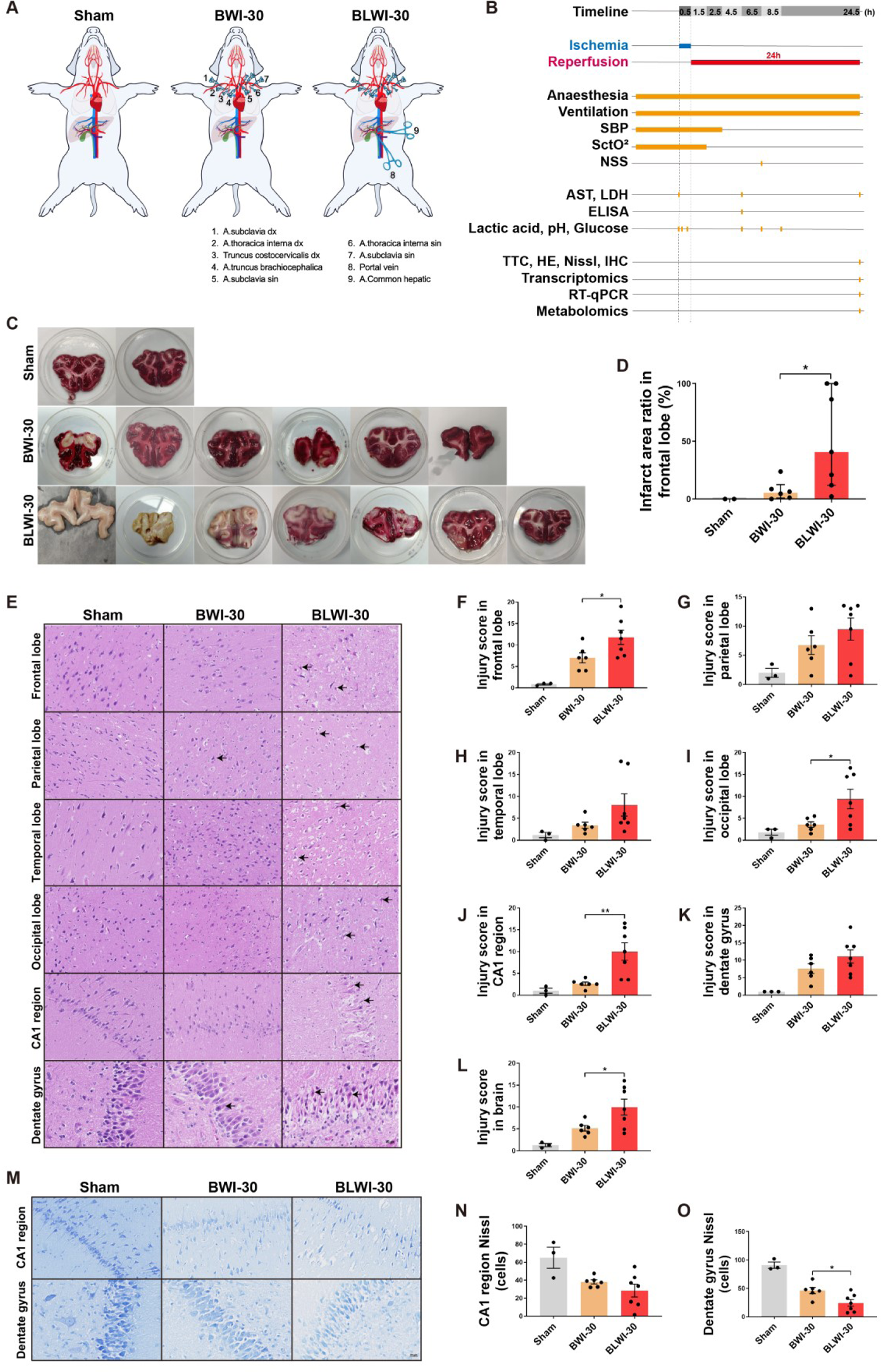
More severe brain injuries in the BLWI-30 versus BWI-30 group. **A**, Schematic of *in vivo* model of the Sham, BLWI-30 and BWI-30 groups. **B**, Timeline of experimental workflow. **C**, Triphenyl-tetrazolium chloride (TTC) staining for the frontal lobe. The white area indicates infarct tissue. **D**, The infarct area ratio in the frontal lobe of three groups. Sham, *N* = 2; BWI-30, *N* = 6; BLWI-30, *N* = 7; two-tailed ratio Mann-Whitney test; histogram, median with interquartile range. **E**, Hematoxylin and eosin staining of the frontal lobe, parietal lobe, temporal lobe, occipital lobe, CA1 region of hippocampus (100×), and dentate gyrus of hippocampus (200×). The arrowhead pointed to the red neuron. **F** to **L**, The injury scores for mean of two fields in the frontal lobe, parietal lobe, temporal lobe, occipital lobe, CA1 region of the hippocampus, dentate gyrus of the hippocampus and the brain (average score of the frontal lobe, parietal lobe, temporal lobe, occipital lobe and hippocampus). **M**, Nissl staining in the CA1 region (100×) and dentate gyrus (200×) of the hippocampus. **N, O**, Live neuron count of the two regions. **F** to **L, N**, and **O**, Sham, *N* = 3; BWI-30, *N* = 6; BLWI-30, *N* = 7; Mean ± SEM, two-tailed ratio unpaired *t*-test; \**P*<0.05, \*\**P*<0.01.

To assess how the liver might impact ischemia reperfusion injury of the brains, we conducted TTC staining to evaluate the infarct area of the frontal lobe, HE staining to assess tissue injury score in different areas of the brain, Nissl staining to assess the viability of the neurons, as well as expression of genes *IL6 and TIMP1 to* assess blood-brain barrier (BBB) damages. TTC staining of the frontal lobe showed that the infarct ratio was higher in the BLWI-30 group than in the BWI-30 group (**Figure 1C and 1D**). Similarly, the tissue injury scores in the frontal lobe, occipital lobe, and CA1 region, were higher in the BLWI-30 group than in the BWI-30 group (**Figure 1E-1H**). As expected, the average tissue injury score of the brain (average score of the frontal lobe, parietal lobe, temporal lobe, occipital lobe, and hippocampus) was much higher in the BLWI-30 group than in the BWI-30 group (**Figure 1L**). Hippocampus is more susceptible to ischemic injury than other brain regions^16,17^. The Nissl staining revealed the number of live neurons in dentate gyrus was lower in the BLWI-30 versus BWI-30 group. The number of live neurons in CA1 region was also lower in the BLWI-30 versus BWI-30 group, although the difference is not statistically significant (**Figure 1M-1O**). Furthermore, expression of *IL6* in the frontal lobe tissue was higher in the BLWI-30 versus BWI-30 group (**Figure S2B**). Expression of *TIMP1* and *IL6* (**Figure S2E and S2F, Table S2**) in the temporal lobe tissue were higher in the BLWI-30 versus BWI-30 group. Taken together, these results indicates that the ischemia reperfusion injury of the brain is exacerbated by simultaneous liver ischemia.

### Increased Intravascular CD45+ Immune Cells of Reperfused Brains with Concurrent Liver Ischemia

BBB damages could induce neuroinflammation^16^. Studies have shown increased immune cell infiltration in the brain tissue during ischemia-reperfusion injury^18,19^. To determine whether concurrent liver ischemia can cause an increase in infiltrating immune cells of the brain, we performed HE and immunohistochemical staining. In HE-stained sections of the five brain regions, more nucleated blood cells were found in the vascular lumen in the brain, particularly in the frontal lobe, of the BLWI-30 versus BWI-30 group (**Figure 2A-2G**). Further identification of immune cells by immunohistochemical staining of CD45 showed more CD45+ cells in the brain, particularly in the frontal lobe, temporal lobe, of the BLWI-30 versus BWI-30 group (**Figure 2H-2N**). These results suggest that the ischemic liver might increase the local immune response during brain ischemia-reperfusion injury.

**Figure 2.**
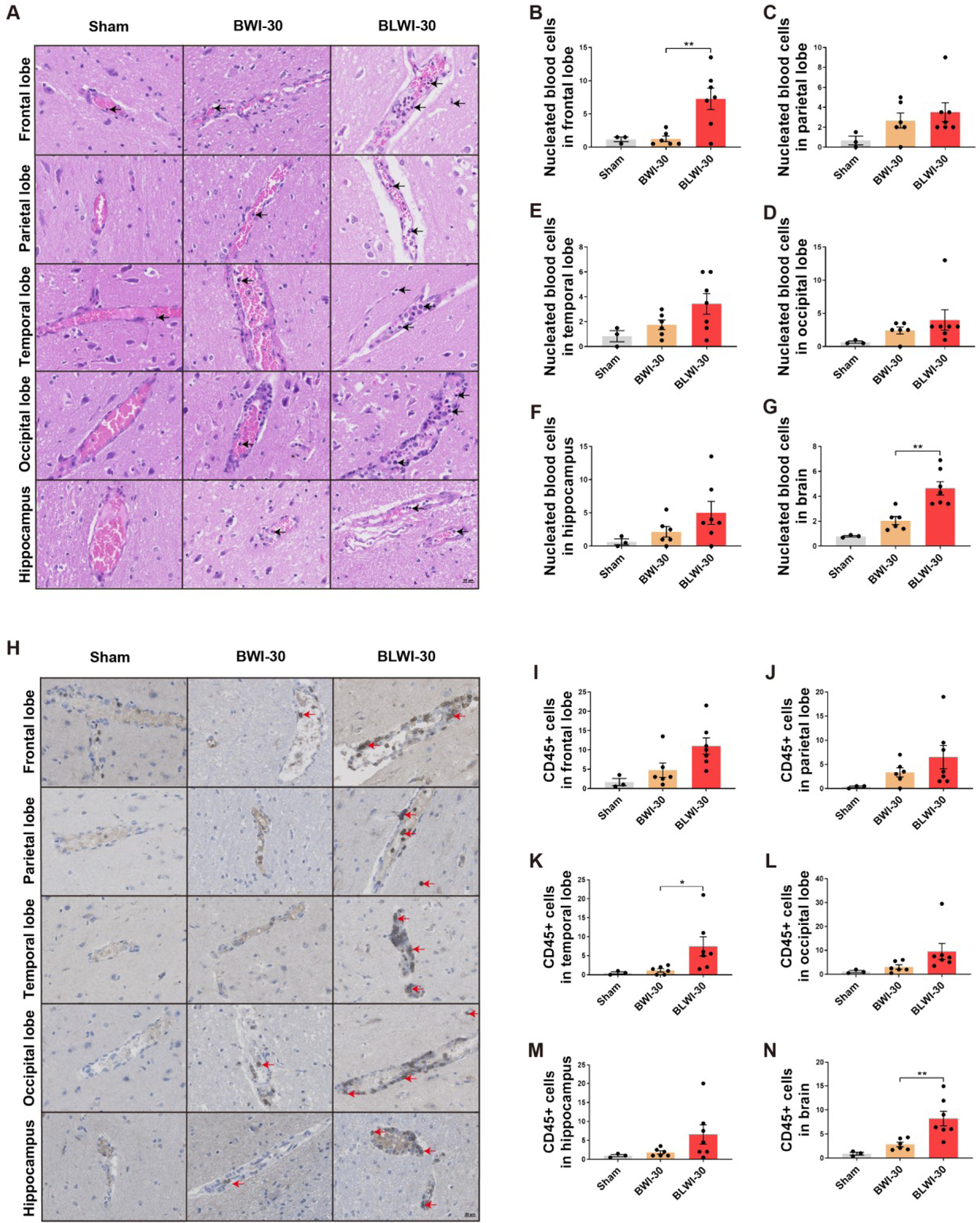
More CD45 positive cells in the BLWI-30 versus BWI-30 *in vivo*. **A**, HE staining showed vascular and nucleated blood cells in the frontal lobe, parietal lobe, temporal lobe, occipital lobe, and hippocampus (200×). The arrowhead pointed to nucleated blood cells. **B** to **G**, The number of nucleated blood cells per field in brain (average number of the frontal lobe, parietal lobe, temporal lobe, occipital lobe, and hippocampus). **H**, Immunofluorescence staining for CD45 in the frontal lobe, parietal lobe, temporal lobe, occipital lobe, and hippocampus (200×). The arrowhead pointed to brown-yellow CD45+ cells. **I** to **N**, The number of CD45+ cells for mean of two fields in the frontal lobe, parietal lobe, temporal lobe, occipital lobe, hippocampus or brain (average number of the frontal lobe, parietal lobe, temporal lobe, occipital lobe and hippocampus). **B** to **G, and I** to **N**, Sham, *N* = 3; BWI-30, *N* = 6; BLWI-30, *N* = 7; Mean ± SEM, two-tailed ratio unpaired *t*-test; \**P*<0.05, \*\**P*<0.01.

**Figure 3.**
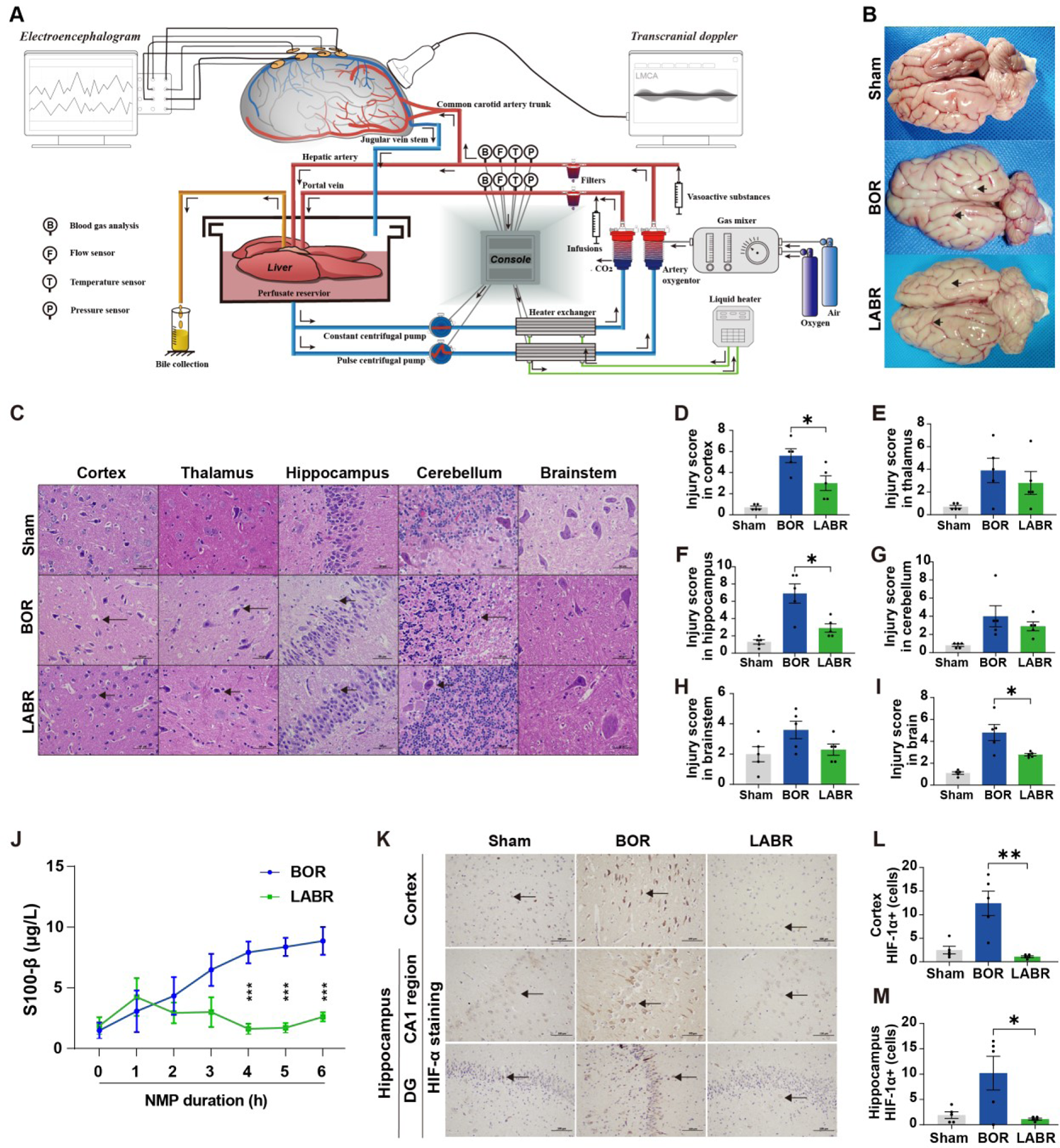
Liver-assisted NMP reduces cell substructure damage and hypoxic injury of the brains. **A**, Technologies for the liver-assisted brain normothermic machine perfusion (NMP) model. **B**, Whole brain structure in the 3 groups. The surface of the brain with cerebral edema, the narrowed sulci (lower arrowhead) and widened gyri (upper arrowhead), in the BOR group. The characteristics were minimal in the LABR group. **C**, Hematoxylin-eosin staining (400×) of cerebral cortex, thalamus, hippocampus, cerebellum, and brainstem after 6 h of NMP (arrowheads pointing to the structures that indicate the neuronal shrinkage after suffering ischemia-reperfusion injury) in the 5 groups. **D** to **I**, The injury scores for mean of two fields in cerebral cortex, thalamus, hippocampus, cerebellum, brainstem or brain (average score of cerebral cortex, thalamus, hippocampus, cerebellum, and brainstem). **J**, The perfusate levels of S100-β. **K**, Immunohistochemistry analysis of hypoxia inducible factor-1α (HIF-1α) indicating the degree of hypoxic injury in the hippocampus and cortex. **L, M,** Number of cells with upregulated expression of HIF-1α were counted in the cerebral cortex (**L**) and hippocampus (**M**). **D** to **I, J, L, M,** *N* = 5, Mean ± SEM; **D** to **I, L, M,** two-tailed ratio unpaired *t*-test; **J**, 2-way ANOVA; \**P*<0.05, \*\**P*<0.01, \*\*\**P*<0.001.

### Restored Brain Circulation and Metabolic Activity after Cardiac Arrest during *ex vivo* Brain NMP

In the *in vivo* global cerebral ischemia model, the clamping of the portal vein transiently affect the hemodynamic stability (**Figure S3A**) and serum lactate level (**Figure S3B and S3C**), which might influence the brain injury. Recently, an *ex vivo* brain NMP model has been developed to investigate pathogenesis of brain diseases^20^. To accurately assess how the liver impacts the recovery of post-CA brain injury, we thus constructed the *ex vivo* brain-only and liver-assisted brain NMP models after cardiac arrest, in which the brain suffered complete instead of global ischemia before reperfusion and hemodynamic stability was not affected by surgical manipulation of the liver. The pressure-controlled NMP provided normothermic, oxygenated perfusion to the brain with whole blood based perfusate to mimic the return of spontaneous circulation after cardiac arrest. The perfusion flow, metabolic activity and electrocortical activity were monitored during *ex vivo* NMP. **Figure 3A, Figure S4A** and **Table S1** demonstrates the technology for the liver-assisted brain NMP.

Due to the preparation of whole blood-based perfusate, the brains in the BOR group underwent a WIT of 10.1 ± 1.8 minutes (**Figure S4A**). Because of the additional procedures, which included removing the liver and connecting it to the system, the mean WIT of the brains was about four minutes longer in the LABR group (14.2 ± 0.7 min) than in the BOR group (*P*=0.0670) (**Figure S4A**). The livers continued to produce bile during perfusion with a portal flow higher than 500 mL/minutes, and with a homogeneous appearance with soft consistency of the parenchyma, indicating that the livers were functioning well. The circulation and oxygen consumption were restored *ex vivo* in both BOR and LABR group. There was no significant difference in the perfusion pressure, pO_2_, pCO_2_, pH, lactate and glucose levels between BOR and LABR during *ex vivo* NMP (**Figure S4B-S4G**). However, the left MCA flow declined in the BOR group, while it maintained stable in the LABR group (**Figure S4H**). Consistently, the arterial resistance declined during the first 2.5 hours but increased thereafter in the BOR group. In contrast, the arterial resistance maintained stable in the LABR group (**Figure S4I**).

Collectively, we have established both *ex vivo* brain only NMP and liver-assisted brain NMP models, during which the circulation and metabolism activity of the brains can be restored *ex vivo* after cardiac arrest.

### Liver-assisted Brain NMP Reduces post-CA Brain Injury

We then tested how addition of the liver in the *ex vivo* NMP circuit might affect the recovery of post-CA brain injury. In line with the increased arterial arterial resistance and reduced perfusion flow, obvious edema of the reperfused brains were observed after 6-h NMP in the BOR group, while edema was not obvious in the LABR (**Figure 3B**). To assess ischemic injuries of the brains, HE staining of different brain areas was conducted and tissue injury score was calculated. There were marked pathological changes including shrunken cells, pyknotic nuclei, and expanded perinuclear space in cortical and hippocampal cells in the BOR group compared to those in the Sham group (**Figure 3C**). In contrast, in the LABR, the pyramidal neurons had plump cell bodies with large, central nuclei and abundant nerve fibers, which was comparable to those in the Sham group (**Figure 3C**). The tissue injury scores in the brains, particularly in the cortex and hippocampus, were lower in the LABR group than in the BOR group (**Figure 3D-3I**).

In addition, the liver-assisted perfused brains had a substantially decreased perfusate level of S100-β, a biomarker of neural injury^21^, compared to the brains in the BOR group (**Figure 3J**). Moreover, immunohistochemistry of hypoxia-induced markers, hypoxia inducible factor-1α (HIF-1α) and heat shock protein 70 (HSP70), antioxygen nuclear factor erythroid 2 like 2 (NRF2) and DNA repairing 8-oxoguanine DNA glycosylase (OGG) in the cerebral cortex was conducted to assess the ischemia-reperfusion injury. The results demonstrated that the brains in the BOR group had a higher density of these markers compared to those in the Sham and LABR groups (**Figure 3K-3M and Figure S5**), suggesting that ischemia-reperfusion injury of the brain is aggravated when the liver is lacking in the NMP circuit. Taken together, the results showed that the liver can protect the brain from post-CA ischemia-reperfusion injury.

### Liver-assisted Brain NMP Improves Neuronal Viability and Protects Cytoarchitecture

Considering that the concurrent ischemia of the liver increased the infarct area of the frontal lobe in the *in vivo* global cerebral ischemia model, we then assessed the density of live neuron by Nissl staining in the *ex vivo* model. The staining revealed comparable densities of live neurons between the Sham group and LABR group in both CA1 region and dentate gyrus. In contrast, the density of live neurons in the CA1 region was lower in the BOR group than in the LABR group (**Figure 4A-4C**). Immunofluorescence analysis of a microglial marker ionized calcium binding adapter molecule 1 (IBA1) produced fragmented signals, with signs of cellular destruction in the CA1 region in the BOR brains but preserved density in the dentate gyrus of all perfused brains in the LABR group (**Figure S6A-S6C**). Staining for an astrocytic marker glial fibrillary acidic protein (GFAP) revealed preserved astrocyte density in all these groups (**Figure S6D-S6F**). Collectively, addition of a liver to the *ex vivo* brain NMP can protect the reperfused brains from cell deaths in the neurons and microglia of the CA1 region.

**Figure 4.**
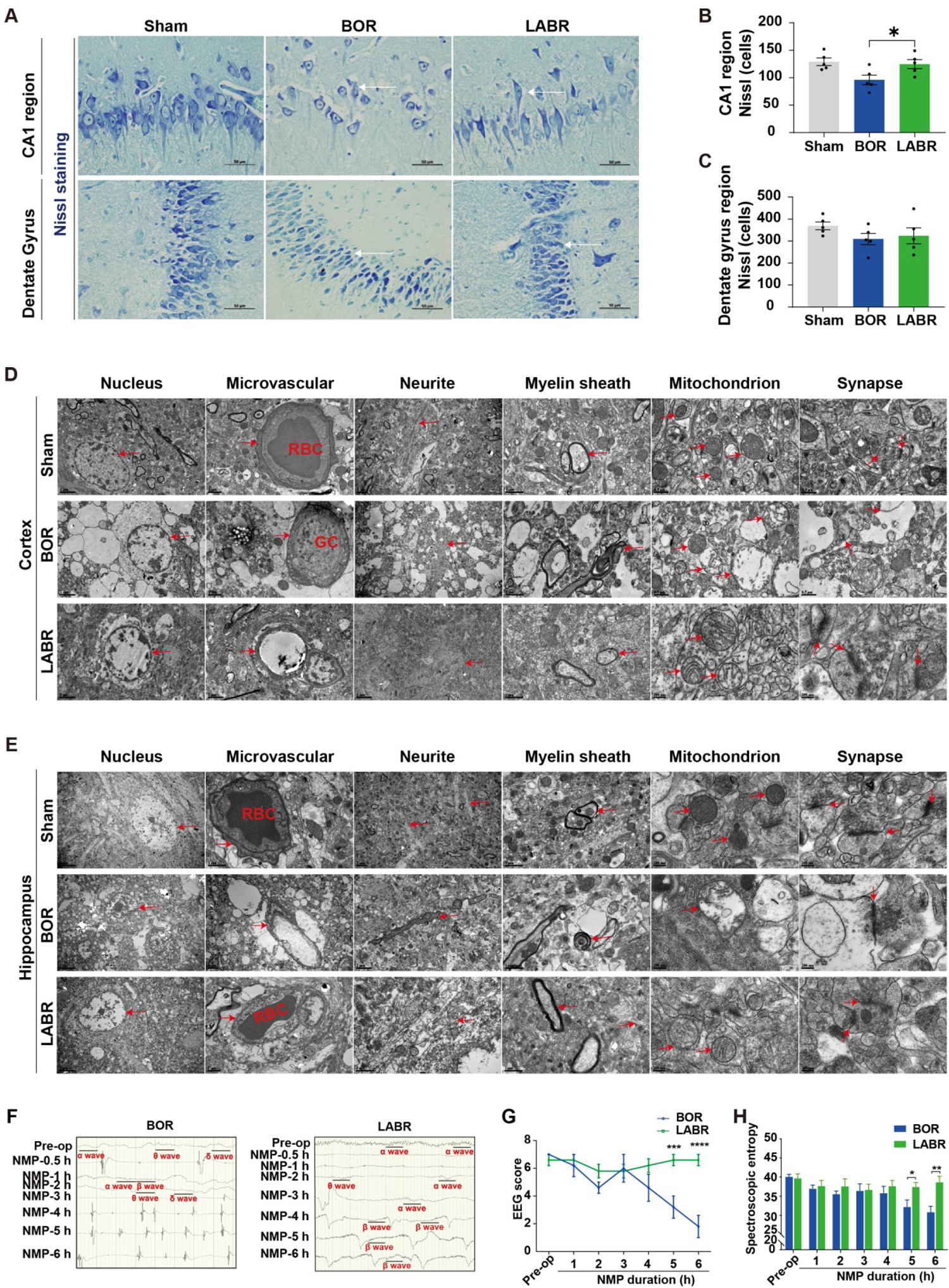
Improved neuronal viability by *ex vivo* liver-assisted brain NMP versus brain only NMP. **A**, Nissl staining (400×) shows the structural integrity of neuronal somas and axons in the hippocampus. **B, C**, Neurons with intact cell bodies were counted in the CA1 region (**B**) and dentate gyrus (**C**). **D, E**, Transmission electron micrographs of the 3 groups in cerebral cortex (**D**) and hippocampus (**E**). The arrowhead pointed to microvessels, neurites, myelin sheaths, mitochondria or synapses. RBC, red blood cell; GC, granulocyte. **F**, EEG results of a representative brain of the BOR or LABR. **G, H**, The EEG score (**G**) and spectroscopic entropy (**H**). **B, G, H**, N = 5, Mean ± SEM; **B, H**, Two-tailed ratio unpaired *t*-test. **G,** 2-way ANOVA; \**P*<0.05, \*\**P*<0.01, \*\*\**P*<0.001, \*\*\*\**P*≤0.0001.

To further evaluate the ultrastructural characteristics of reperfused brains, we conducted the TEM analysis of the brain tissues. In the cerebral cortex and hippocampus of the BOR group, severe tissue vacuolization, mild microvascular edema, light loose myelin sheath, massive mitochondrial swelling or membrane rupture, mitochondrial cristae rupture and degradation, and significant synaptic degradation, were documented. In contrast, in the majority of cells of the LABR group, the microvessels, neurites, myelin sheaths, mitochondria, synapses, and neurotransmitters were structurally intact. Erythrocytes were observed in the microvascular lumen of the LABR and Sham groups, whereas granulocytes were observed in the BOR group (**Figure 4D and 4E**), which is in line with *in vivo* studies that more immune cells trapped in the vessels of the brains with concurrent liver ischemia. Collectively, addition of a liver in the *ex vivo* brain NMP circuit, can improve neuronal viability and cytoarchitecture.

### Liver-assisted Brain NMP Maintains Global Electrophysiological Activity

It is of relevance to detect whether the reduced brain injury can be translated to improved global electrophysiological activity in the LABR group. The EEG monitoring showed that the α or (and) β waves, both of which are considered to represent conscious activity^22,23^, presented within 30 minutes after the start of NMP but disappeared after 3-5 h of NMP in four of five brains in the BOR group (**Figure 4F and Table S3**). In contrast, frequent α and β waves showed up within 1 h of NMP and maintained until the end of 6-hour NMP in all the perfused brains in the LABR group (**Figure 4F and Table S3**). The EEG score and spectroscopic entropy were much higher at the end of NMP in the LABR group than in the BOR group (**Figure 4G and 4H**). Collectively, these data show that electrocortical activity can be maintained *ex vivo* by liver-assisted brain NMP instead of brain-only NMP.

To determine the longest period of warm ischemia after which global electrophysiological activity can be restored and maintained *ex vivo* in pigs, we extended the warm ischemia time of the brains to 30 minutes (LABWI-30), 50 minutes (LABWI-50), 60 minutes (LABWI-60), or 240 minutes (LABWI-240) under the support of a liver (**Figure S7**). The global electrophysiological activity was restored in all five brains, and the presence of α or β wave was maintained until the end of NMP in four of five brains in the LABWI-30 group and in all five brains in the LABWI-50 group (**Table S3**). In the LABWI-60 group, all the brains presented with α or β wave during 3-4 h of NMP, although the waves disappeared soon (**Table S3**). In the LABWI-240 group, no α or β wave was observed (**Table S3**). These results indicate that the global electrophysiological activity of the brain can be restored and maintained *ex vivo* after a warm ischemia of over 50 minutes, when a functioning liver is added in the NMP circuit.

### Transcriptome Analysis Reveals a Reduced Cell Death and Immune Response in Brain Tissues without versus with Simultaneous Hepatic Ischemia

To elucidate the molecular mechanisms underlying the protective effects of the liver on brain injury, RNA-seq was employed to profile the frontal and temporal lobes of the brain with or without concurrent hepatic ischemia. In the frontal lobe, we observed significant up-regulation and down-regulation of genes in the presence of simultaneous hepatic ischemia. Specifically, we identified 245 up-regulated genes and 350 down-regulated genes in the brain with simultaneous hepatic ischemia (BLWI-30 group) (**Figure 5A**). Functional annotation of the differentially expressed genes highlighted distinct functional terms. The up-regulated genes in the BLWI group were enriched in GO terms related to the regulation of programmed cell death and immune response, such as positive regulation of apoptotic process, regulation of T cell chemotaxis, and B cell activation (**Figure 5B**). Conversely, the down-regulated genes in BLWI-30 were enriched in GO terms associated with neuronal functions, including neuron projection development and regulation of neurotransmitter levels (**Figure 5B**). Similar transcriptome differences were observed in the temporal lobe, with 269 up-regulated genes and 337 down-regulated genes in the BLWI-30 group (**Figure 5C**). These differential genes exhibited comparable functional gene ontology term enrichments (**Figure 5D**). These molecular alterations corroborate our prior findings on brain function and pathological analyses, indicating that the support of a functioning liver can reduce brain damage caused by ischemia. Furthermore, among the down-regulated genes, we also identified differential genes involved in metabolic processes. For instance, genes (*AK7*, *AK9*, *TYMS*) involved in nucleotide metabolism may be a consequence of cell death and immune response (**Figure S8A and S8B**).

**Figure 5.**
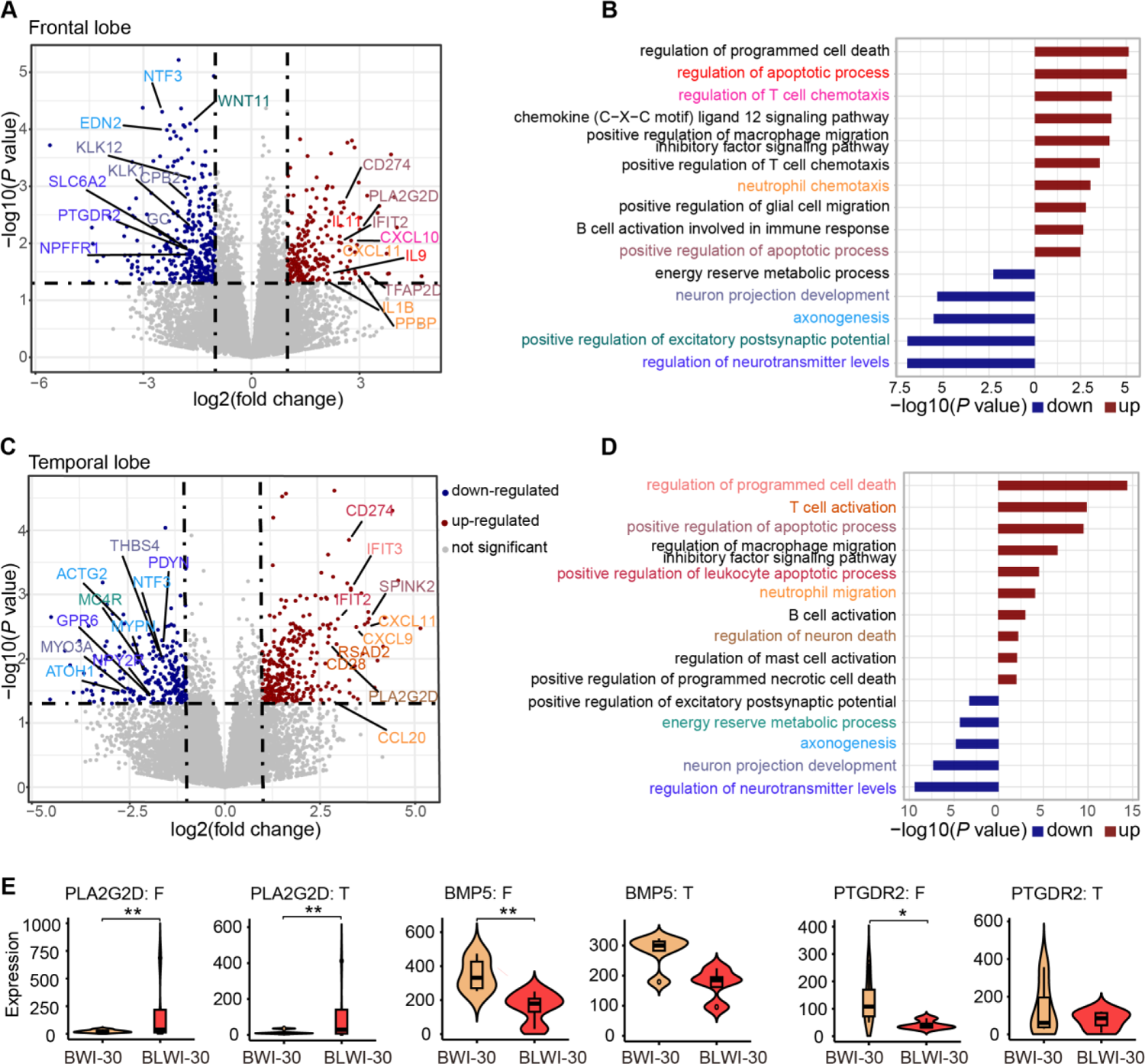
Transcriptome Differences between Brain Tissues with and without Simultaneous Hepatic Ischemia by RNA-seq. **A**, Volcano plot displaying the differential genes in the frontal lobe of the brain with (BLWI group) and without (BWI group) simultaneous hepatic ischemia. Genes significantly up-regulated or down-regulated in BLWI group (*P* value <0.05, fold change >1) are represented in red and blue, respectively. Functional interest genes were labeled according to their annotated functional terms as shown in (**B**). **B**, Bar plot illustrating GO enrichment of significant differential genes in the frontal lobe. Up-regulated and down-regulated genes were tested separately and presented in red and blue, respectively. GO terms were color-coded corresponding to (**A**). **C**, Volcano plot depicting the differential genes in the temporal lobe of the brains of the two groups. Genes significantly up-regulated or down-regulated in BLWI group (*P* value <0.05, fold change >1) are represented in red and blue, respectively. Functional interest genes were labeled according to their annotated functional terms as shown in (**D**). **D**, Bar plot displaying GO enrichment of significant differential genes in the temporal lobe. Up-regulated and down-regulated genes were tested separately and presented in red and blue, respectively. GO terms were color-coded corresponding to (**C**). **E**, Violin plot showing gene expression (RPKM) in each biological replicate (N=4). \**P*<0.05, \*\**P*<0.01.

Intriguingly, we also found differential genes enriched in terms of response to energy reserve metabolic processes and found genes involved in lipid, linoleic (*PLA2G2D*), and ketone (*BMP5*, *PTGDR2*) metabolism processes (**Figure 5E**), as well as in glycolysis/gluconeogenesis (*ALDOB*/*PCK1*) and GAD in aspartate and glutamate metabolism, taurine, and hypotaurine metabolism (**Figure S8C and S8D**).

The results collectively underscore the intricate interplay between hepatic function and brain tissue physiology, highlighting the systemic influence of hepatic ischemia on brain gene expression and metabolism.

### Dramatic Metabolic Differences in Brain Tissues with and without Simultaneous Hepatic Ischemia

To unveil the metabolic dynamics during brain ischemia, especially the difference between brain ischemia with or without liver ischemia, we further performed UHPLC-QTOFMS analysis of the brain frontal lobe and temporal lobe regions. The supervised partial least squares analysis (PLS-DA) of the overall metabolic profiles from 5-6 biological replicates in the two groups clearly demonstrated distinct metabolism differences (**Figure 6A and 6B, Figure S9A and S9B**). The distribution of metabolism classes in general, was summarized in **Figure S9C** and **S9D**.

**Figure 6.**
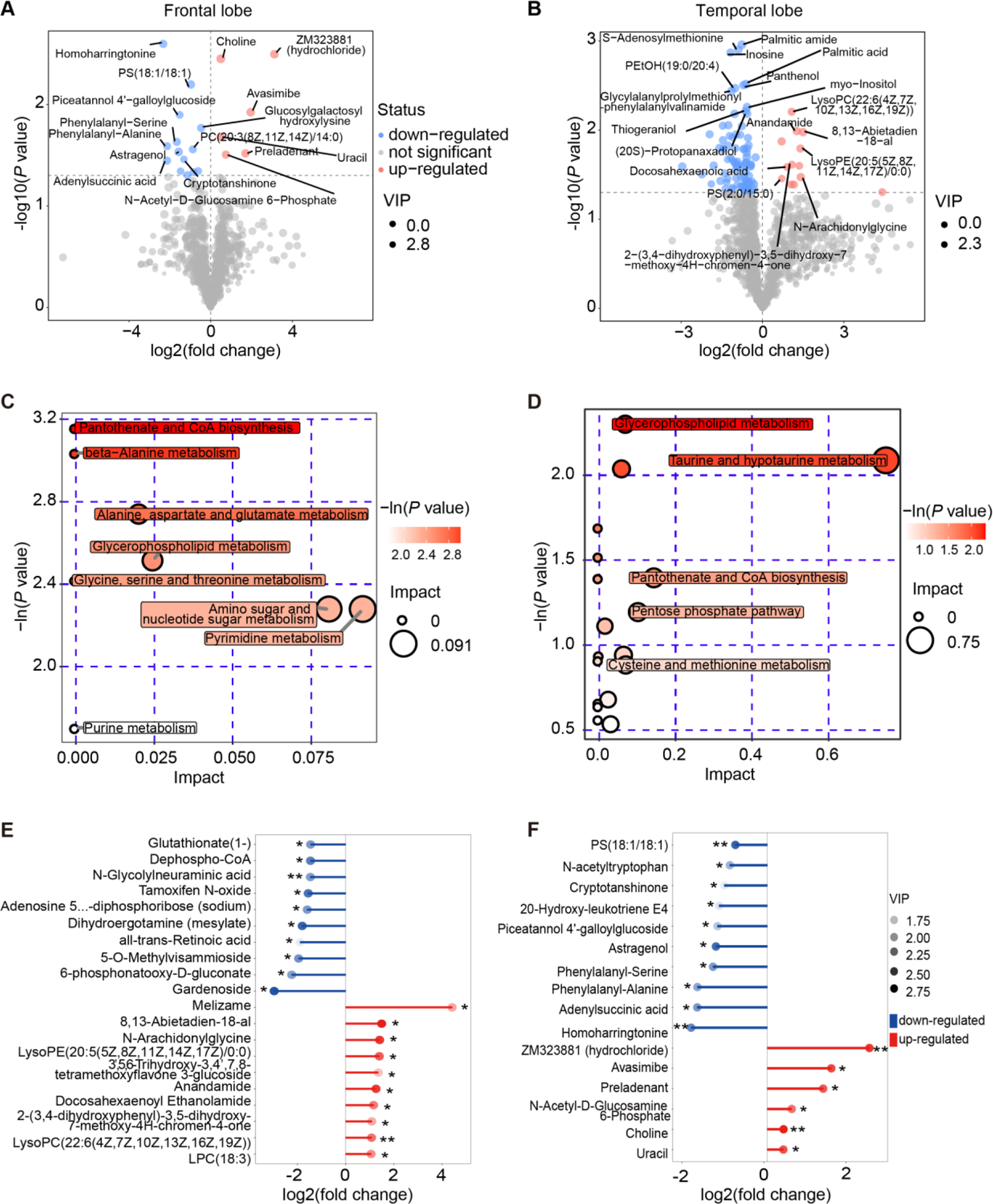
Metabolic Differences in Brain Tissues with and without Simultaneous Hepatic Ischemia by UHPLC-QTOFMS. **A, B**, Volcano plot illustrating the differential metabolites in the frontal lobe (**A**) and temporal lobe **(B**) region of the brain with and without simultaneous hepatic ischemia. Significant metabolic alterations (*P* value<0.05, fold change >1) were denoted in red or blue, respectively. **C, D**, Bubble plot displaying the enriched metabolic pathways for increased metabolites for frontal lobe and temporal lobe respectively. The value on the x-axis and the size of the circle indicates the impact of the difference. The value on the y-axis and the color indicates the significance of enrichment in -ln(*P*). **E, F**, Matchstick plot illustrating the top metabolic alterations. The x-axis displays the log2(fold change) of BLWI to BWI. Each item’s VIP score is denoted by color.

Interestingly, we found that the down-regulated metabolism in the BLWI-30 group (higher in the BWI-30 only group) was significantly enriched in pantothenate and CoA biosynthesis, glycerophospholipid metabolism pathways in both brain regions (**Figure 6C and 6D**). Further analysis revealed that metabolites in the linoleic acid, arachidonic acid, and glycerophospholipid pathways, are decreased in the BLWI-30 group (**Figure S8E and S8F**). Both linoleic acid and arachidonic acid metabolism are crucial energy metabolism pathways and is important for brain function. Linoleic acid, arachidonic acid and their derivatives are essential components in the modulation of synaptic transmission and neurotransmitter release, thereby influencing various neurological processes^24^. Linoleic acid, arachidonic acid and their derivatives may play a role in promoting neuroprotection and supporting neuronal survival under various pathological conditions, including ischemic injuries^25^. In the opposite, the up-regulated metabolism in the BLWI-30 group was fewer and enriched in divergent pathways such as, pantothenate and CoA biosynthesis, beta-Alanine metabolism, Cholinergic synapse (**Figure S9E** and **S9F**). The most significantly different metabolite was shown in **Figure 6E** and **7F**, including variant compounds contributing energy metabolism and neuronal functions.

**Figure 7.**
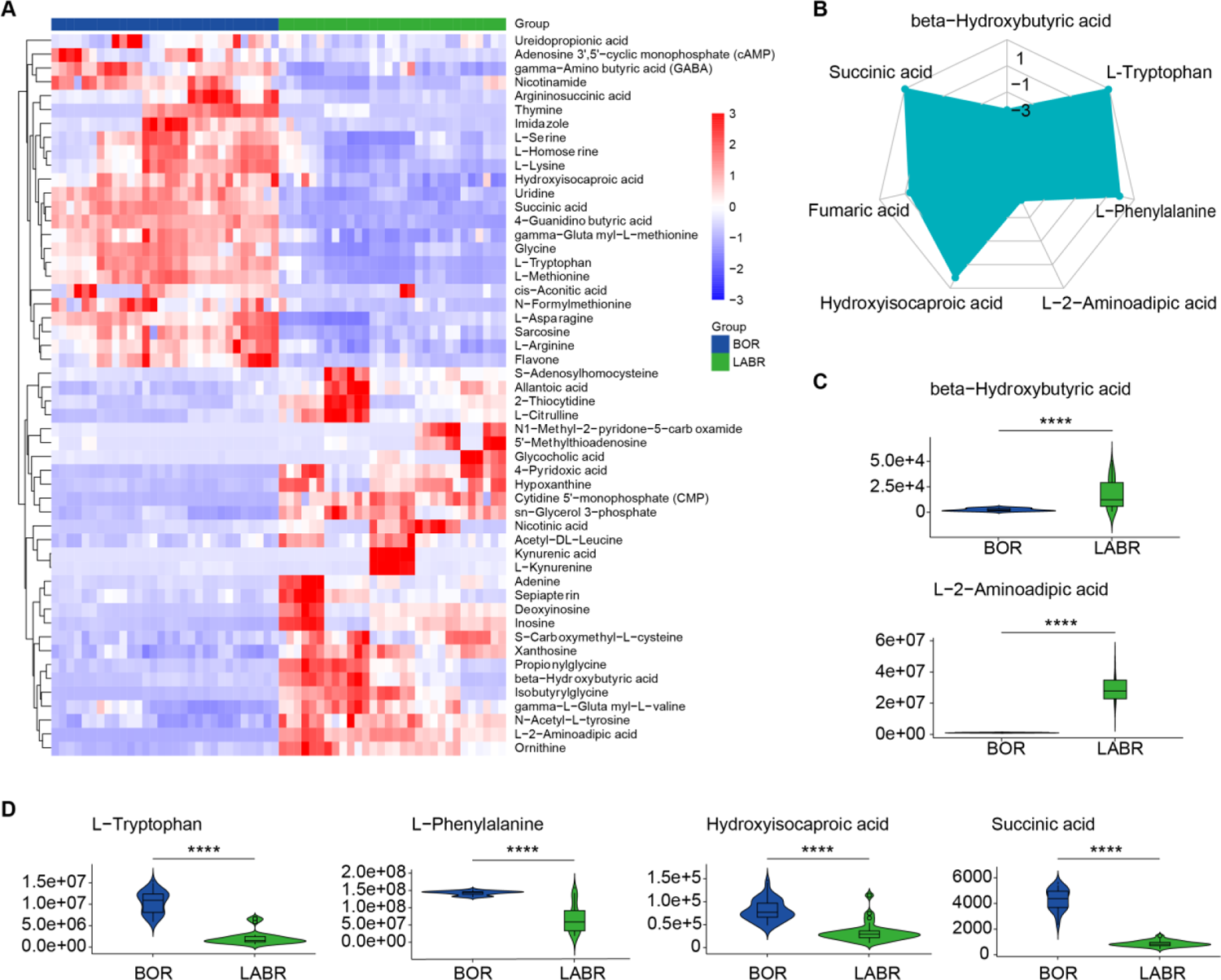
Metabolic Differences in the Perfusate Serum during *ex vivo* Brain Normothermic Machine Perfusion with and without the Support of a Functioning Liver. A, Heatmap displaying significant different metabolites, presented by z-score across each row. **B**, Radar chart illustrating the differences by metabolic groups. **C**, Violin plot showing the signal for decreased metabolites in *ex vivo* normothermic machine perfusion without the support of a functioning liver (decreased in the BOR group). **D**, Violin plot showing the signal for increased metabolites in *ex vivo* normothermic machine perfusion without the support of a functioning liver (increased in the BOR group). **C, D**, Two-tailed ratio unpaired *t*-test; \*\*\*\**P*≤0.0001.

The observed differential gene expressions and metabolic shifts in the presence or absence of hepatic ischemia highlight the critical influence of the liver on brain metabolism and gene expression during ischemic conditions. These findings provide additional evidence on the molecular level, supporting the notion that the liver plays a significant role in protecting brain function, particularly under ischemic conditions.

### Metabolic Differences of Perfusate during *ex vivo* Brain NMP with and without Support of a Functioning Liver

The analysis of the metabolome difference in our global cerebral ischemia model with or without liver ischemia revealed a dramatic difference in metabolism. This difference was contributed by multiple dimensions, including both energy metabolism differences as well as the response to cell deaths and immune activation. To illustrate how the liver may contribute to the energy metabolism of the brain, we further conducted ultra-high-performance liquid chromatography coupled to triple-quadrupole mass spectrometry (UHPLC-QqQ-MS) to reveal the perfusate metabolomic profiles in the BOR and LABR groups after 2-3 hours of NMP. Both the OPLS–DA analysis (**Figure S10A and S10B**) and differential analysis showed different metabolomic profiles between the two groups (**Figure S10C and S10D**).

The hierarchical clustering analysis showed that there are 39 differential metabolites (VIP > 1 and *P*<0.05) between BOR versus LABR (**Figure 7A and Figure S10E**). Interestingly, we found that β-hydroxybutyrate acid (ketone body) was the most significantly decreased metabolism in the absent of liver (BOR group). While most of the upstream or intermedia metabolites in ketone body metabolism, L-Phenylalanine, L-Tryptophan, and Hydroxyisocaproic acid were significantly decreased in the present of liver (LABR group) (Figure 7B-7D). In situations where glucose availability is limited, the liver produces ketone bodies, such as β-hydroxybutyrate and acetoacetate, through the process of ketogenesis^26^. These ketone bodies serve as alternative energy substrates for the brain, particularly during periods of fasting. These results indicates that during *ex vivo* Brain NMP, the liver may support the brain via enhanced energy metabolism, including production of ketone bodies^26^.

With our simple *ex vivo* system, we found a significant metabolic difference in the perfusate, indicating that metabolism mediates organ communication. The results collectively underscore the intricate interplay between hepatic function and brain tissue physiology, highlighting the systemic influence of hepatic ischemia on brain metabolism and gene expression.

## DISCUSSION

Post-CA brain injury accounts for two thirds of total deaths in patients with CA who survived the first 48-72 h^27^. Clinical studies have shown that both pre-CA cirrhosis and post-CA hypoxic liver injury are associated with significantly poorer survival and neurological outcomes in patients with CA^11,12,28,29^. To the best of our knowledge, we show for the first time the direct evidences that the liver plays a crucial role in pathogenesis of post-CA brain injury.

The pathophysiology of post-CA brain injury is encompassed by initial (ischemic) injury and secondary (reperfusion) injury^27^. The major presentations of ischemic injury at the cellular level is cessation of aerobic metabolism with consequent depletion of adenosine triphosphate (ATP), which subsequently leads to massive infux of sodium and water and intracellular cytotoxic oedema. Intracellular Ca^2+^ overload and activation of the innate immunity and subsequent tissue inflammation are the two components of the reperfusion injury. In this study, we found in the *in vivo* isolated global cerebral ischemia model that the ratios of infarction in the frontal lobe were larger, in the brains with concurrent hepatic ischemia than those without. Moreover, the tissue injury was more severe with increased intravascular immune cell adhesion in the brains with concurrent hepatic ischemia, particularly in the CA1 region. In consistent with the *in vivo* data, we found that lack of a liver in the *ex vivo* brain NMP circuit reduced cytoarchitectural integrity and electrocortical activity of the brains in comparison to those with liver support. Collectively, both the *in vivo* and *ex vivo* results show that the liver can profoundly protect post-CA brain injury.

In addition, our studies on both the transcriptome and metabolome levels provide significant insights into the molecular mechanisms underlying the protective effects of the liver on post-CA brain injury. Our findings indicate that the absence of a functioning liver during ischemic conditions leads to the up-regulation of genes associated with cell death and immune response in the brain. Conversely, the down-regulation of genes related to neuronal functions, such as neuron projection development and regulation of neurotransmitter levels. These alterations likely contribute to the reduction of neuronal damage and preservation of synaptic function under the protective effects of the liver. These molecular alterations are in consistent with our functional observation and pathophysiological assays at cellular level.

Notably, the identification of differentially expressed genes associated with the energy reserve metabolic process underscores the potential role of liver-mediated metabolic regulation in mitigating the detrimental effects of ischemia on brain tissue. This suggests a possible mechanism through which the liver’s metabolic regulatory functions could contribute to the overall protection of the brain during ischemic insults. Previous studies have demonstrated that hepatic soluble epoxide hydrolase (sEH) activity was specifically altered following traumatic brain injury (TBI) and negatively correlated with the plasma levels of 14,15,-epoxyeicosatrienoic acid, which shows neuroprotective effects in the controlled cortical injury mouse model^30^. Hepatic sEH activity also plays roles in regulating cerebral Aβ metabolism and the pathogenesis of Alzheimer’s disease in mice^31^. Further investigation to identify and validate the exact molecules involved in the protection of the brain during ischemic insults is required and will be of great clinical relevance for advancing therapeutic interventions.

In clinical practice, various advanced life support technologies have been used to resuscitate patients with CA, including electricdefibrillation, mechanical ventilation, extracorporeal membrane oxygenation and continuous renal replacement therapy^32^. The results of the current study showed profound pathological, corticoelectral and metabolic impacts of the liver on post-CA brain injury. However, currently no liver replacement therapy has been used during CA resuscitation. Considering the complex function of the liver, targeting limited genes, proteins or metabolites might not be an ideal method. Artificial extracorporeal liver support (AELS) might be a feasible option to this end^33^, although the efficacy and duration of AELS are still suboptimal^34^. Extracorporeal support with pig or human liver perfusion^35,36^, might also be a potential therapeutic option. Recent progress in gene-edited minipigs have made clinical xenotransplantation possible^37–40^. These minipigs can provide a timely organ supply for extracorporeal liver support. Moreover, the techniques for long term (7-14 days) *ex situ* liver NMP have recently been developed^41,42^. Therefore, the feasibility and efficacy of extracorporeal support with liver perfusion might be enhanced and translated into clinical practice in the field of post-CA brain injury.

Undoubtedly, there were limitations in this study. Firstly, due to the complex function of the liver and the weakness of mechanism studies in pigs, we could not identify specific genes, proteins, metabolites or immune cells which can fully explain the protection of the liver on the brain, although our data suggest that ketone bodies might be an important target against post-CA brain injury. Subsequently, no therapeutic strategy to enhance the liver function for brain protection has been tested in the current study. Finally, the results of *ex vivo* electrophysiological activity assessment suggest that the brain might tolerate 30-50 minutes WIT, which is far beyond the common belief of the maximum ischemic tolerance (5-8 minutes) of the brain. However, the *ex vivo* assessment of brain function are limited. Comprehensive assessment of whole brain function is required. These limitations provide room for future studies.

In conclusion, the results from the current study showed the crucial role of the liver in the pathogenesis of post-cardiac arrest brain injury. These findings shed lights on a novel cardio-pulmonary-hepatic-cerebral resuscitation strategy. Moreover, the *ex vivo* liver-assisted brain NMP model provides a unique platform for further investigating the maximum ischemic tolerance of the brain, and the roles of other organs in post-cardiac arrest brain injury. The knowledge from the current and future studies potentially improves survival and outcomes of patients suffering CA.

## Sources of Funding

This study was supported by grants as follows: the National Natural Science Foundation of China (81970564, 82170663, 82370664, to Dr Guo; 82070670, to Dr He; 82300744, to Dr Xu), the Guangdong Provincial Key Laboratory Construction Projection on Organ Donation and Transplant Immunology (2023B1212060020) (to Dr He), Guangdong Provincial International Cooperation Base of Science and Technology (Organ Transplantation) (2020A0505020003) (to Dr He), Science and Technology Program of Guangdong (2020B1111140003) (to Dr He). This study was also supported by China organ transplantation development foundation (to Dr Guo) and Medical Scientific Research Foundation of Guangdong Province of China (A2020275 to Dr Yin).

## Disclosures

None.

## Supplemental Material

### Methods

Tables S1–S3

Figure S1–S10

## Notes

### Competing Interest Statement

The authors have declared no competing interest.

